# A toolbox for nodule development studies in chickpea: a hairy-root transformation protocol and an efficient laboratory strain of *Mesorhizobium* sp

**DOI:** 10.1101/362947

**Authors:** Drishti Mandal, Senjuti Sinharoy

## Abstract

*Mesorhizobium sp*. produces root nodules in chickpea. Chickpea and model legume *Medicago truncatula* are members of inverted repeat lacking clade (IRLC). The rhizobia after internalization inside plant cell called ‘bacteroid’. Nodule Specific Cysteine-rich (NCR) peptides in IRLC legumes guide bacteroids to a ‘terminally differentiated swollen (TDS)’ form. Bacteroids in chickpea are less TDS than those in *Medicago*. Nodule development in chickpea indicates recent evolutionary diversification and merits further study. A hairy root transformation protocol and an efficient laboratory strain are prerequisites for performing any genetic study on nodulation. We have standardized a protocol for composite plant generation in chickpea with a transformation frequency above 50%, as shown by fluorescent markers. This protocol also works well in different ecotypes of chickpea. Localization of subcellular markers in these transformed roots is similar to *Medicago*. When checked inside transformed nodules, peroxisomes were concentrated along the periphery of the nodules, while ER and golgi bodies surrounded the symbiosomes. Different *Mesorhizobium* strains were evaluated for their ability to initiate nodule development, and efficiency of nitrogen fixation. Inoculation with different strains resulted in different shapes of TDS bacteroids with variable nitrogen fixation. Our study provides a toolbox to study nodule development in the crop legume chickpea.

## Introduction

Root nodule symbiosis (RNS) is the most successful metabolism-dependent symbiosis on earth. Leguminous plants get reduced nitrogen directly from RNS, at the expense of photosynthate (Werner et al., 2015). The staple crop chickpea (*Cicer arietinum*) is world’s second largest cultivated legume. Chickpea seeds are a valuable source of dietary protein in lower socio-economic class. Additionally, due to the symbiotic interaction with *Mesorhizobium sp*., chickpea can be cultivated in a sustainable way (Jain et al., 2013; Varshney et al., 2013). Research on nodule development has been centered upon model legumes *Medicago truncatula* and *Lotus japonicus.* Today, model legumes are in the forefront of legume biology in terms of both available knowledge and resources. The only disadvantage is that model legumes are not crop species. In spite of being the most important grain legume in tropical and sub-tropical countries (Jukanti et al., 2012), literature on chickpea nodule development is scarce. Genome sequencing and the establishment of transcriptomic and proteomic resources have laid the pillars for making chickpea a model amongst crop legumes (Jain et al., 2013; Varshney et al., 2013; Ramalingam et al., 2015; Pandey et al., 2018).

Chickpea belongs to the inverted repeat lacking clade (IRLC), which diverged from model legume *Medicago* ~10-20 million years ago (Jain et al., 2013; Varshney et al., 2013). IRLC legumes develop indeterminate nodules, with a gradient of cells at different stages of development can be seen from the distal to the proximal part of the nodule. A persistent meristem (zone I) is present at the distal end of the nodule. Bacterial endocytosis and colonization takes place in the postmeristematic cells of the infection zone (zone II), where plant membrane-bound bacterial units are formed. These units are considered as an ammonium-exporting organelle called symbiosome (Roth and Stacey, 1989). The rhizobia inside the symbiosomes are called bacteroids. The bacteroids divide inside the symbiosome and gradually colonize the whole cell. In the nitrogen fixation zone (zone III) bacteroids differentiation is terminated, ammonium assimilation genes are repressed, and nitrogen fixation genes are induced (Oldroyd, 2013; Udvardi and Poole, 2013).

Terminally differentiated, enlarged bacteroids with different morphotypes are a typical feature of the IRLC legumes. The major determining factor behind these irreversibly differentiated bacteroids is nodule specific cysteine-rich (NCR) peptides (Montiel et al., 2016; Montiel et al., 2017). The molecular mechanism of NCR peptide mechanism of NCR peptide regulated-endo-reduplication of the symbiont genome has been worked out in model legume *Medicago. Medicago* genome encodes more than 700 *NCR* genes. At least 138 NCR peptides get processed in the endoplasmic reticulum (ER) and targeted towards symbiosomes (Wang et al., 2010; Durgo et al., 2015). NCR peptides force endo-reduplication of the symbiont genome. As a result, the bacteria (now bacteroid) lose their ability to divide and re-grow on culture media, but their size increases up to ~10-fold (Mergaert et al., 2006; Young et al., 2011; Sinharoy et al., 2013). *NCR* gene family evolved in *Medicago* by a recent local gene duplication (Alunni et al., 2007). *NCR*s have been identified in several IRLC legumes, including chickpea (Kant et al., 2016; Montiel et al., 2016; Montiel et al., 2017). The number of *NCR* genes among different IRLC legumes varies greatly. This results in different morphotypes of bacteroids in different legumes, such as swollen/spherical, elongated, and elongated-branched (Montiel et al., 2017). *Mesorhizobium TAL620*-induced nodules in chickpea express only 63 *NCR* genes, while its’ bacteroids endo-reduplicated up to 4 fold (Kant et al., 2016; Montiel et al., 2017). Chickpea and *Medicago* NCRs share less than 80% identity. Chickpea NCR peptides have more identity with *Glycyrrhiza uralensis, Onobrychis vicifolia,* and *Astragalus canadensis*, while phylogenetically chickpea is closer to *Medicago* (Montiel et al., 2016; Montiel et al., 2017). Chickpea’s swollen/ spherical bacteroids are basal to the evolution of NCR-guided morphogenesis (Montiel et al., 2017). In contrast to chickpea, *Medicago* has more than 700 *NCR* genes produce elongated-branched bacteroids depicting an advanced stage of this trait. Interestingly, this morphogenesis of bacteroids and the genome endo-reduplication is thought to determine the efficiency of nitrogen fixation in respective legumes (Oono and Denison, 2010). Thus, comparative investigation of nodule development in chickpea and *Medicago* will enhance our knowledge on the evolutionary link between variable nitrogen fixation efficiencies amongst different legumes.

An efficient hairy-root transformation protocol is an indispensable tool to understand nodule biology, enabling us to study the localization of any protein (fluorescent tag), activities of promoters (GUS fusion), and the effects of over-expression or knock-down of certain genes. We have undertaken an effort to establish hairy root transformation in chickpea and study nodule development. Unlike *A. tumefaciens, A. rhizogenes* generates transformed roots from the site of infection. *A. rhizogenes* contains *root locus* (*rol*) genes in the Ri-plasmid, which promotes the formation of genetically transformed adventitious hairy roots. *A. rhizogenes* carrying a recombinant Ri plasmid can generate composite plants. These plants contain untransformed shoot and transform root (Georgiev et al., 2012). When such an *A. rhizogenes* additionally carrying a gene of interest in a binary vector is used for transformation, a certain percentage of co-transformed roots are obtained. Both overexpression and downregulation of a specific gene can be achieved by these co-transformed roots (Limpens et al., 2004; Sinharoy and DasGupta, 2009; Sinharoy et al., 2015). Recently, it has been reported that *CRISPR/Cas9* mediated gene knock out is also possible in hairy roots (Ron et al., 2014; Cai et al., 2015; Wang et al., 2016). Till date, all the successful protocol for hairy root generation in other legumes have shown the transformed roots to be biologically similar to untransformed roots, with no difference in normal nodule development (Stougaard et al., 1987; Quandt et al., 1993; Stiller et al., 1997; Boisson-Dernier et al., 2001; Van-de-Velde et al., 2003; Limpens et al., 2004; Estrada-Navarrete et al., 2006; Kereszt et al., 2007; Sinharoy et al., 2009; Bonaldi et al., 2010; Imanishi et al., 2011; Brijwal and Tamta, 2015; Thilip et al., 2015; Habibi et al., 2016; Thwe et al., 2016). *A. rhizogenes*-mediated hairy-root transformation is an efficient and less time-consuming alternative method for the functional validation of genes. Here we are reporting an efficient protocol for hairy root transformation in chickpea. We have developed both *in vivo* and *ex vitro* protocols. We have used four different *Mesorhizobium* strains to follow nodule initiation and the developmental program and evaluated the nitrogen fixation efficiency. We have also checked the localization of several subcellular markers in transformed chickpea roots and nodules. Additionally, we have confirmed that the shape of the bacteroids in the mature nitrogen-fixing zone of chickpea nodules is directly correlated with the efficiency of nitrogen fixation.

## Results

### Hairy root transformation of chickpea

We used *Agrobacterium rhizogenes R1000, ARqua1*, and *MSU440* strains for hairy root transformation. All three strains were successful to generate hairy roots (data not shown). For detailed characterization, we used *ARqua1* strain. Dicotyledonous plants can be transformed using both *in vivo* (tissue culture based), and *ex vitro* (without tissue culture) protocols (Collier et al., 2005; Sinharoy et al., 2015). We attempted both the methods. For *in vivo* transformation, chickpea seedlings were infected either by cutting 1-3 cm of the radicle and scraping it in an *A. rhizogenes* lawn culture or using a needle containing *A. rhizogenes* to make a small wound at the hypocotyl region. Overview of the methods is given in Figure 1. In both the cases, transformed roots began to emerge from the infected region 14-15 days after infection (dai) (Figure 1 step-Ia). In case of needle mediated *rhizogenes* infection, the original roots were removed from the plant when transformed root started coming. These plants were transferred to kanamycin containing selection medium to increase the efficiency of the transformation. We tried two different concentration of kanamycin (50µg/ml, and 25µg/ml). The plants were maintained in the antibiotic-containing medium for 10-12 days for antibiotic selection. All the roots became black, and the plant died within two weeks when 50 µg/ml of kanamycin was used (data not shown). But the transgenic roots grew normally in presence of 25 µg/ml kanamycin. The shoots of composite plants had no difference in morphology with their non-transgenic counterparts (Figure 1 step IIa). The transformed roots exhibited characteristics of plagiotropic growth. The transformation frequency was determined using either cauliflower mosaic virus (*CaMV*) *35S* promoter, or *Arabidopsis* ubiquitin 10 (*AtUBQ10*) promoter-driven expression of the DsRed fluorescent protein (Figure 1 step IIIa). Both showed a similar frequency of transformed roots (Table 1). Usually, we obtained a slightly greater frequency of transformed roots after antibiotic selection (Table 1. 48.7% without antibiotic over 53.1% with an antibiotic in *BDG256*). Plants generated by *in vivo* methods were transferred to the substrate (a mixer of 3:1 vermiculite and fireclay balls) 20-25 dai. Non-fluorescent roots were removed from the plant at this point. Plants were maintained in this substrate for further 2-3 days under nitrogen free condition. Nodules were observed in these transformed roots after 10 days post-inoculation (dpi) with *Mesorhizobium* (Figure 1 Step IVa).

**Figure 1:**
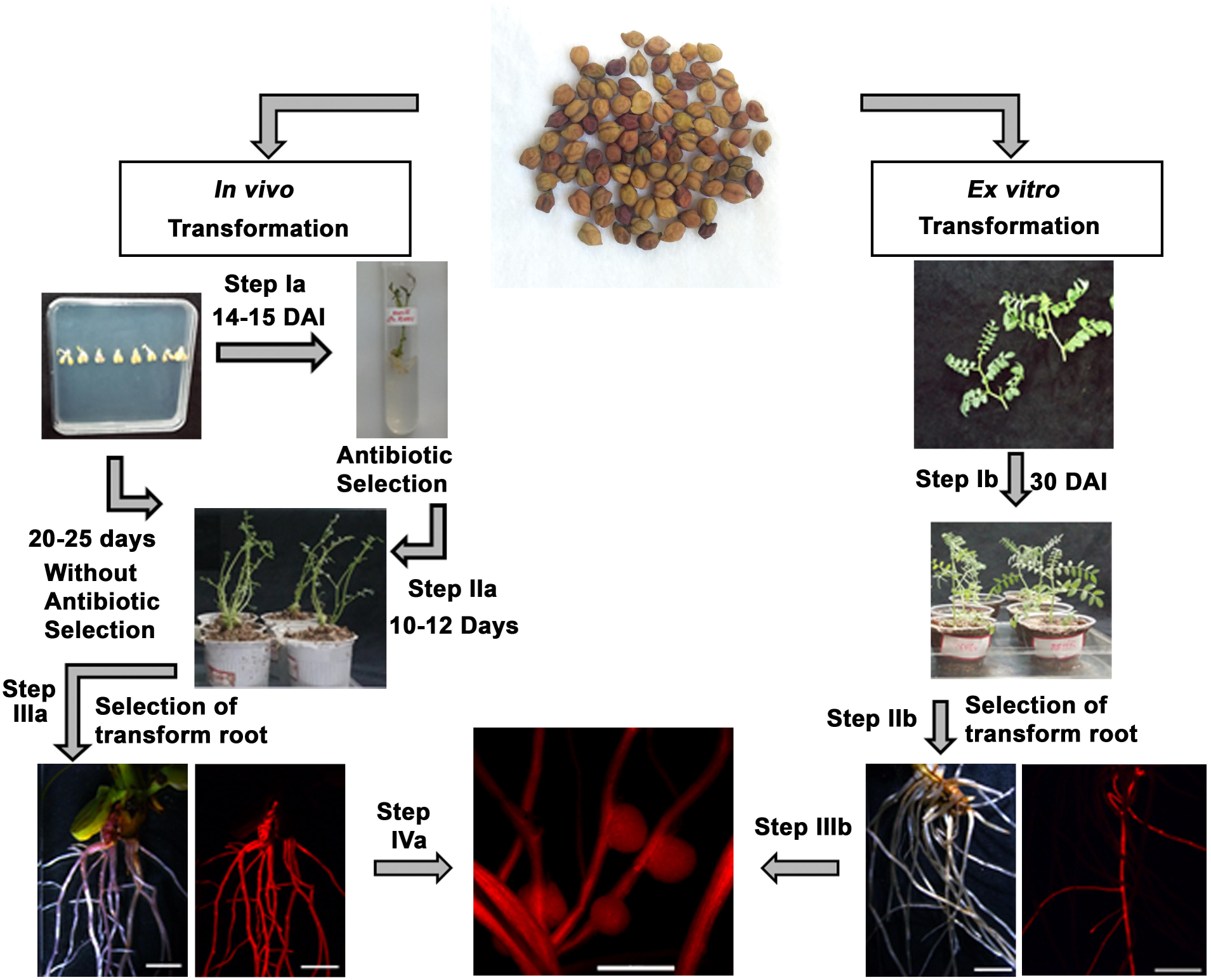
*Agrobacterium rhizogenes* induced hairy roots in chickpea. *In vivo* transformation: Germinated seedlings were infected with *Agrobacterium rhizogenes* and plated on Fahraeus medium. 14-15 days after infection hairy roots were seen and the composite plants were transferred in selective antibiotics (depicted in step Ia). After 10- 12 days, the plants were transferred to growing substrate (depicted in step IIa). Transformed roots were screen under the red channel of the stereo-microscope (depicted in step IIIa). Composite plants were inoculated with *M. ciceri CC1192* after 10 days (depicted in step IVa). *Ex vitro* transformation: The wounded portion was inoculated with *Agrobacterium rhizogenes.* 30 days after infection the hairy roots were seen (depicted in step Ib) and transformed roots were screened under the red channel of stereo-microscope (depicted in step IIb). The composite plants were inoculated with *M. ciceri CC1192* for nodulation (depicted in step IIIb). The transformed root pictures were taken under stereo-microscope either under bright field or under red channel.

**Table 1:** *Agrobacterium rhizogenes* mediated transformation of different chickpea ecotypes with *pKGW-Red-Root*:

| Genotype | Region | Method of transformation |  | Efficiency of transformation |  |
| --- | --- | --- | --- | --- | --- |
|  |  | *In vivo* | *Ex vitro* | *In vivo* | *Ex vitro* |

|  |  | Cut | Inject |  | Cut | Inject |  |
| --- | --- | --- | --- | --- | --- | --- | --- |
| ICC4958<br>(DESI) | India | 20(16) | ND | ND | 26.027% | ND | ND |
| BDG256<br>(DESI) | India | 20(19) | 20(17) | 20(14) | 48.717% | 39.614% | 38.333% |
| BDG256<br>(DESI) | India | 25(21) | ND | ND | 53.135%* | ND | ND |
| ICC17258<br>(DESI) | India | 20(20) | ND | ND | 40.331% | ND | ND |
| ICC1885<br>(DESI) | India | 20(18) | ND | ND | 37.313% | ND | ND |
| ICC8261<br>(KABULI) | Turkey | 20(9) | ND | ND | 26.984% | ND | ND |
| Local Variety<br>(DESI) | India | 35(35) | 35(31) | 35(34) | 50.259%* | 51.437%* | 40.107% |

\* % after antibiotic selection

*Ex-vitro* composite plant generation is a rapid, efficient, simple, and low-cost method for producing composite plants without tissue culture (Collier et al., 2005; Sinharoy et al., 2015). The steps for *ex-vivo* transformation are shown on the right-hand side of Figure 1. For *ex-vitro* transformation, 25-30 days old apical stem tissue of young branches were cut and dipped in the *A. rhizogenes* culture and transferred to the substrate (Figure 1 Step1b). Further, the plants were maintained in a humid chamber for 30 days to promote root production in presence of full nitrogen fertilization regime. Both transformed and adventitious root production were observed by 20 dai. The transformed roots were screened based on dsRed fluorescence (Figure 1 step IIb). At this point, the substrate containing plants were washed thoroughly with distilled water, and transferred to a new pot and maintained under nitrogen free condition. After 2-3 days the plants were inoculated with *Mesorhizobium* culture for root nodule development, and nodules were noticed within 10 dpi (Figure 1 step IIIb).

We have tested this protocol in different chickpea ecotypes. A list of different cultivars of chickpea from different origin, where we successfully tested our transformation protocol, has been given in Table 1. Both *in vivo* and *ex vitro* methods had comparable transformation efficiency. While we obtained greater transformation efficiency in *BDG256*, and ‘local’ variety (above 50%), *ICC4958, ICC8261*, and kabuli had a lower efficiency. In summary, this method can be used in several different varieties of chickpea cultivars.

### Expression of different subcellular markers in transformed hairy roots and nodules of chickpea

To test the activity of transgenes transformed following our protocol we employed already known markers for their subcellular localization in chickpea root and nodules. These markers were already published for either *Arabidopsis thaliana* (*At*) (model plant), or *Medicago truncatula* (*Mt*) (model legume) (Nelson et al., 2007; Ivanov and Harrison, 2014). We expressed some of these markers in hairy roots and nodules of chickpea. A construct created by fusing the signal peptide of *AtWAK2* (*Arabidopsis thaliana* wall-associated kinase 2) at the N-terminus of a *mCherry* sequence which also contained endoplasmic reticulum (ER) retention signal HDEL at the C-terminus, was used as ER marker. This *mCherry* was under *AtUBQ10* promoter (He et al., 1999). An extensive characteristic tubular and sheet-like ER network was seen throughout the cytoplasm of chickpea root epidermal cells (Supplemental Figure 1A). As expected, ER network around the nucleus was more uniform (Supplemental Figure 1A). The mitochondrial marker was created by adding the first 29 amino acids of *Saccharomyces cerevisiae* cytochrome c oxidase IV with a *mCherry* sequence under *pAtUBQ10* (Kohler et al., 1997). Typical small and round mitochondria were present in high copy numbers in every cell in chickpea hairy root epidermis. The mitochondria were generally clustered in the periphery of the cells, and around the nucleus (Supplemental Figure 1B). A similar result was obtained earlier in *Medicago* (Ivanov and Harrison, 2014). We have used *pAtUBQ10* driven *mCherry* fused with microtubule binding domain of mammalian microtubule-associated protein 4 (Marc et al., 1998) as the microtubule marker. In root epidermal cells, we noticed extensive microtubular networks (Supplemental Figure 1C). The marker for peroxisome was created by fusing SKL signal peptide of *AtPTS1* to the C-terminal of *mCherry* (Reumann, 2004). Innumerable peroxisomes were seen in the cells of chickpea root epidermis (Supplemental Figure 1D). Unlike mitochondria, peroxisomes were scattered all around the cells (compare Supplemental Figure 1B and D). To monitor the localization of apoplasts, we used the *pAtUBQ10*-driven expression of *Medicago* blue copper protein’s (*MtBCP1*) 23 amino acid signal peptide (*BCPsp*) fused to *mCherry* (Pumplin and Harrison, 2009). MtBCP1 signal peptide drives the routing of mCherry to apoplasts, as was also seen in *Medicago* before (Supplemental Figure 1E) (Ivanov and Harrison, 2014). From the *MtBCP1sp-mCherry* localization, it is clear that the epidermal cells are connected through apoplastic connection (Supplemental Figure 1E). Additionally, we used *p35S* promoter driven expression of a plasma membrane aquaporin (*PIP2a*) fused to cyan fluorescent protein (*CFP*) (Cutler et al., 2000). We noticed uniform labeling of CFP along the surface of the cell (Supplemental Figure 1F). A marker created by fusing plasmodesmata-located protein 1 (*PDLP1*) with *mCherry* at N terminus under *pAtUBQ10* was used to visualize the cytoplasmic connections between the cells (Thompson and Wolniak, 2008). A cross-section of chickpea root showed the sym-plastic connections between the cortical cells (Supplemental Figure 1G). The golgi body marker was created by fusing N-terminal 49 amino acids of *GmMAN1* to the N terminus of *mCherry* (Saint-Jore-Dupas et al., 2006). The golgi apparatus were scattered throughout the cytoplasm (Supplemental Figure 1H) in the chickpea root epidermal cell. To check the hormone responsiveness in these transformed roots, we used a construct created by fusing a nuclear localization signal (*NLS*) at the C-terminus of *GFP* under auxin responsive synthetic *DR5* promoter (*DR5-GFP-NLS*) (Suzaki et al., 2013). We noticed high *GFP* expression around the proliferating lateral root primordial cells (Supplemental Figure 1I), but GFP fluorescence was not observed in the mature root cortex (Supplemental Figure 1I). In summary, by using eight different subcellular markers under both *35S* and *AtUBQ10* promoter, and the *DR5-GFP* construct, we confirmed the normal physiological functioning of these transformed roots (Supplemental Figure 1).

The transgenic hairy roots also showed efficient nodule development (Figure 1). After the bacterial endocytosis in the post-mitotic cells, bacteroids multiply and rapidly fill up the entire cell. The peribacteroid membrane (PBM) is a plant-derived membrane. Hence, the division of bacteroids accompanies a huge expansion of the elements of host cell membrane. Several parallel evidences suggest that membrane trafficking towards symbiosomes increases significantly during this phase (Gavrin et al., 2017). Symbiosomes in *Medicago* has a unique mosaic identity with plasma membrane syntaxin SYP132, and endomembrane marker (regulatory small GTPases of the Rab family) Rab7 (Limpens et al., 2009). This suggests that under certain circumstances, established markers may behave differently inside the infected nodule cells. To demonstrate the utility of this protocol, we used three subcellular markers (peroxisome, ER, and golgi) to check their localization inside nodule cells. Peroxisomes are organelles that play key roles in plant cell metabolism (Hu et al., 2012). The central infected nodule tissue is surrounded by three layers of uninfected peripheral tissues. Namely from inside to outside, the nodule parenchyma, endodermis, and cortex (Xiao et al., 2014). We noticed the presence of a significantly higher number of peroxisomes in the nodule cortex around the vascular tissue (Figure 2A). The number of peroxisomes were significantly low in nodule parenchyma, and endodermis (Figure 2A). But the concentration of peroxisomes were more or less similar in all infected zones (Figure 2B-C). The golgi apparatus (cis-golgi) were noticed everywhere inside the infected cells surrounding the symbiosomes (Figure 2D). We failed to detect any golgi apparatus surrounding the infection thread (Figure 2D). ERs also show similar localization pattern to the golgi, close to the symbiosomes (Figure 2E). Additionally, we were able to detect ERs around the infection thread (Figure 2F).

**Figure 2:**
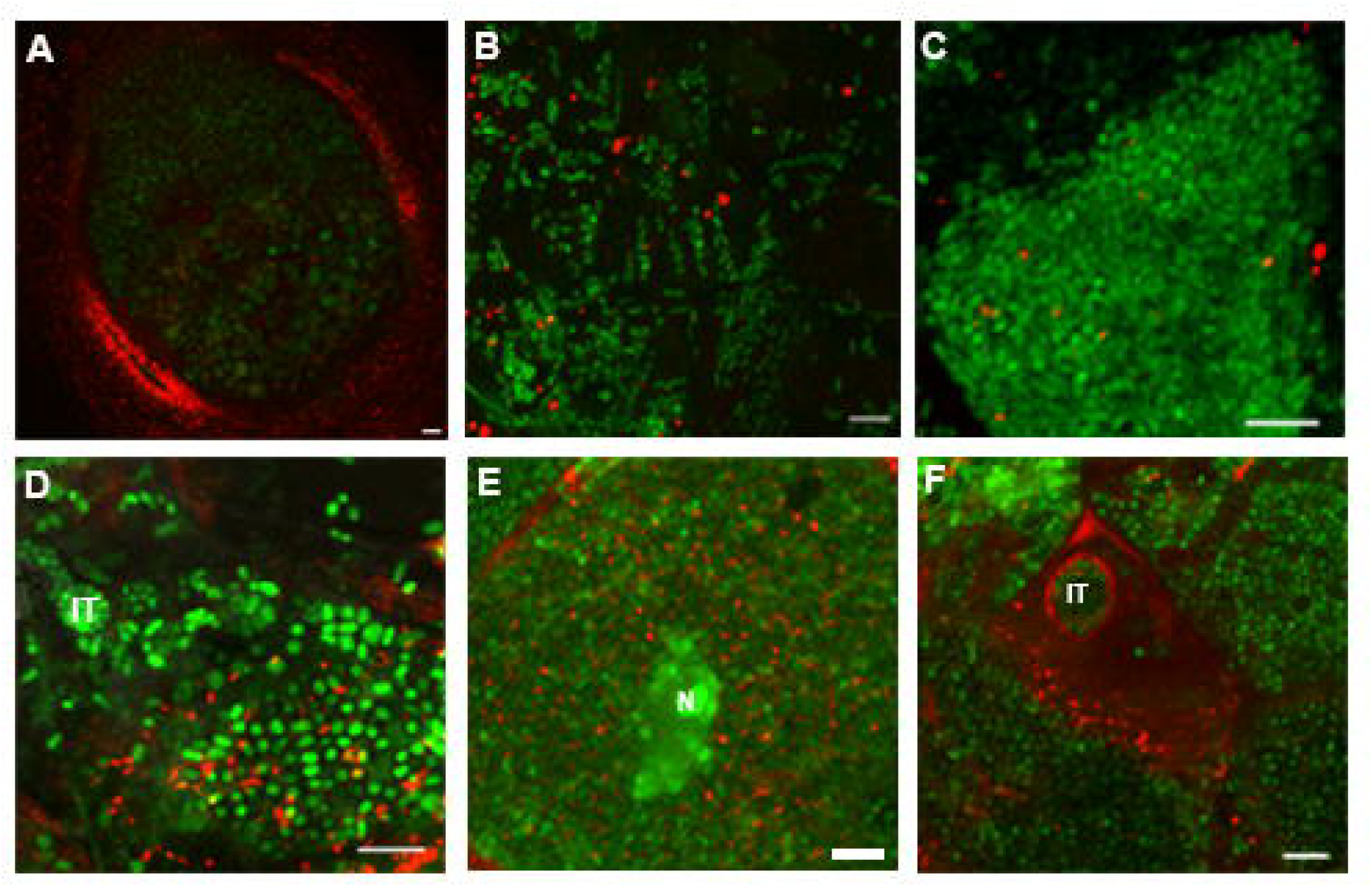
Localization of different subcellular protein markers in transgenic chickpea nodules**: A-F**, Transformed nodules were hand sectioned and stained with SYTO13 (DNA, green). **A-C**, mCherry-SKL showing localization **A**, whole nodule **B**, invasion zone **C**, nitrogen fixation zone **D**, MAN49-mCherry localization in the nitrogen fixation zone **E-F**, mCherry-HDEL localization in endoplasmic reticulum in the nitrogen fixation zone of transgenic chickpea nodules induced by *M. ciceri CC1192*. Nucleus marked by N and infection thread marked by IT. Scale bars in **A**, = 50µm and **B-F**, = 5µm.

### Evaluating the symbiotic potential of different *Mesorhizobium* strains

An efficient laboratory strain is a pre-requisite to study nodule development. We examined several different *M. ciceri* strains originating from different sources to evaluate their potential for healthy nodule development, and N_2_ fixation efficiency. *M. ciceri CC1192* is an Australian isolate used as a commercial inoculant there, and recently the complete genome sequence has been published (Haskett et al., 2016). *M. ciceri IC-59* was isolated by ICRISAT, India (Rupela and Sudarshana, 1990; Esfahani et al., 2014), and being used for different field trials (Esfahani et al., 2014). *M. ciceri TAL620* was used for studies in past (Chandra and Pareek, 1987), and recently has been used for the transcriptome sequencing of chickpea root nodules (Kant et al., 2016). Our own isolation effort from the experimental garden of National Institute of Plant Genome Research (NIPGR), India, has resulted to the culturing of NIPGR Isolate 7 or *NI-7*. Chickpea plants with healthy pink nodules were collected from the experimental garden. The pink nodules were dissected, surface sterilized, and rhizobia were isolated from the nodules (detail in material and method). The most efficient strain (*NI-7*) was used for the comparative study. The *16S rDNA* sequence of the *NI-7* strain has been submitted to NCBI (MH517450). Interestingly, *NI-7* shows the highest similarity with *Mesorhizobium tianshanense. M. tianshanense* nodulating *Cicer arietinum* and *Cicer canariense* have already been reported (Rivas et al., 2007; Armas-Capote et al., 2014). We generated nodules by inoculation of all four above mentioned *Mesorhizobium* strains (Supplemental Figure 2). All four strains were able to initiate nodules by 6 dpi (bump formation) (Supplemental Figure 2 A-D). Numerous small white round shaped nodules were observed by 10 dpi (Supplemental Figure 2 E-H). At 15 dpi, nodules were pink (due to the expression of leghemoglobin) in case of *CC1192, IC59*, and *NI-7* (Supplemental Figure 2 I-K) but remained white in case of *TAL620* (Supplemental Figure 2L). When followed further, all four nodules become pink by 21 dpi (Supplemental Figure 2 M-P). By 28 and 35dpi *CC1192, IC59*, and *NI-7* nodules showed similar developmental fate by becoming elongated and cylindrical (Supplemental Figure 2 Q-S and U-W), while *TAL620* nodules were comparably smaller and pale pink (Supplemental Figure 2 T and X). Senescence zones, a typical feature of indeterminate nodules, started appearing by 35 dpi in most of the nodules (Supplemental Figure 2 U-X). In case of *TAL620* nodules, the senescence zone was visible even from 28 dpi (Supplemental Figure 2 T). This suggests that *TAL620* is a comparably inefficient strain for chickpea nodule development. We performed acetylene reduction assay (ARA) at 21 and 35 dpi to determine the nitrogen fixation efficiency of these strains. *CC1192* showed highest ARA, followed by *NI-7, IC59,* and *TAL620* at both the time points (Figure 3A). *TAL620* showed significantly low ARA compared to *CC1192* (the most efficient strain), at both the time points. *IC59* shows significantly low ARA only at 35 dpi. Taken together, our data show that *CC1192* is the most efficient strain for chickpea followed by *NI-7*. Further, we have used SYTO dyes to stain the nitrogen fixation zone of these chickpea nodules at 21 dpi. Interestingly, we found small spherical symbiosomes inside *CC1192, IC59*, and *NI-7* generated nodules (Figure 3 B-D), whereas in case of *TAL620* the symbiosomes were small and elliptical (Figure 3E). Our result shows a direct correlation of the shape of bacteroids with ARA. Acidification of symbiosome space is crucial for functional nitrogen fixation. LysoTracker DND189 has been successfully used in *Medicago* for *in planta* detection of peribacteriod space (PBS, also called symbiosome space) acidification (Pierre et al., 2013). We then asked the question whether there is any correlation between the PBS acidity and ARA by the above mentioned strains. We used pH sensitive dyes DND189 to stain the most efficient (*CC1192*), and the least efficient (*TAL620*) *Mesorhizobium*-induced nodules. DND189 is a weak base linked with a fluorophore. It gets protonated in acidic organelles and is retained in the membranes of those organelles. The biggest advantage of DND-189 over most of the lysotrackers is, it is almost non-fluorescent except when inside acidic compartments (Lopez et al., 2005). DND189 starts fluorescing at a pH of ≤ 5.2 and reaches an optimal emission between pH 4 and 5. We used DND189 in combination with SYTO82/PI to check the acidity of chickpea symbiosome space. In case of *CC1192*, DND-189 was accumulated in the PBS and we observed the clear spherical shape of the acidic zone (PBS) (Figure 4A-B). The bacterial nucleic acid showed clear red staining when it was counter-stained with SYTO82 (Figure 4A). There was some acidity in the PBS of the symbiosomes made by strain *TAL620*, but we rarely obtained a clear demarcation of PBS boundary when stained with DND189 (Figure 4C). To test the viability of bacteria inside *TAL620*-induced nodules, we used propidium iodide (PI) and SYTO staining (Haag et al., 2011). Live bacteria with intact cytoplasmic membranes would be stained by membrane-permeable SYTO dye, while dead bacteria with degraded cytoplasmic membranes would be stained by membrane-impermeable PI. When *TAL620* induced nodules were stained with PI, the nucleic acid in most of the samples did not take up the stain. Interestingly, the few instances where sporadic symbiosomes took up the stain, they were spherical (Figure 4D). But the frequency of occurrence of such zone inside the nodules is very scare. Whereas SYTO dye shows live bacteroids inside *TAL620* induced nodules (Figure 3E) Hence, from our observation it appears that DND-189 is accumulated less specifically in the PBS of *TAL620* induced symbiosomes, indicative of less acidity.

**Figure 3:**
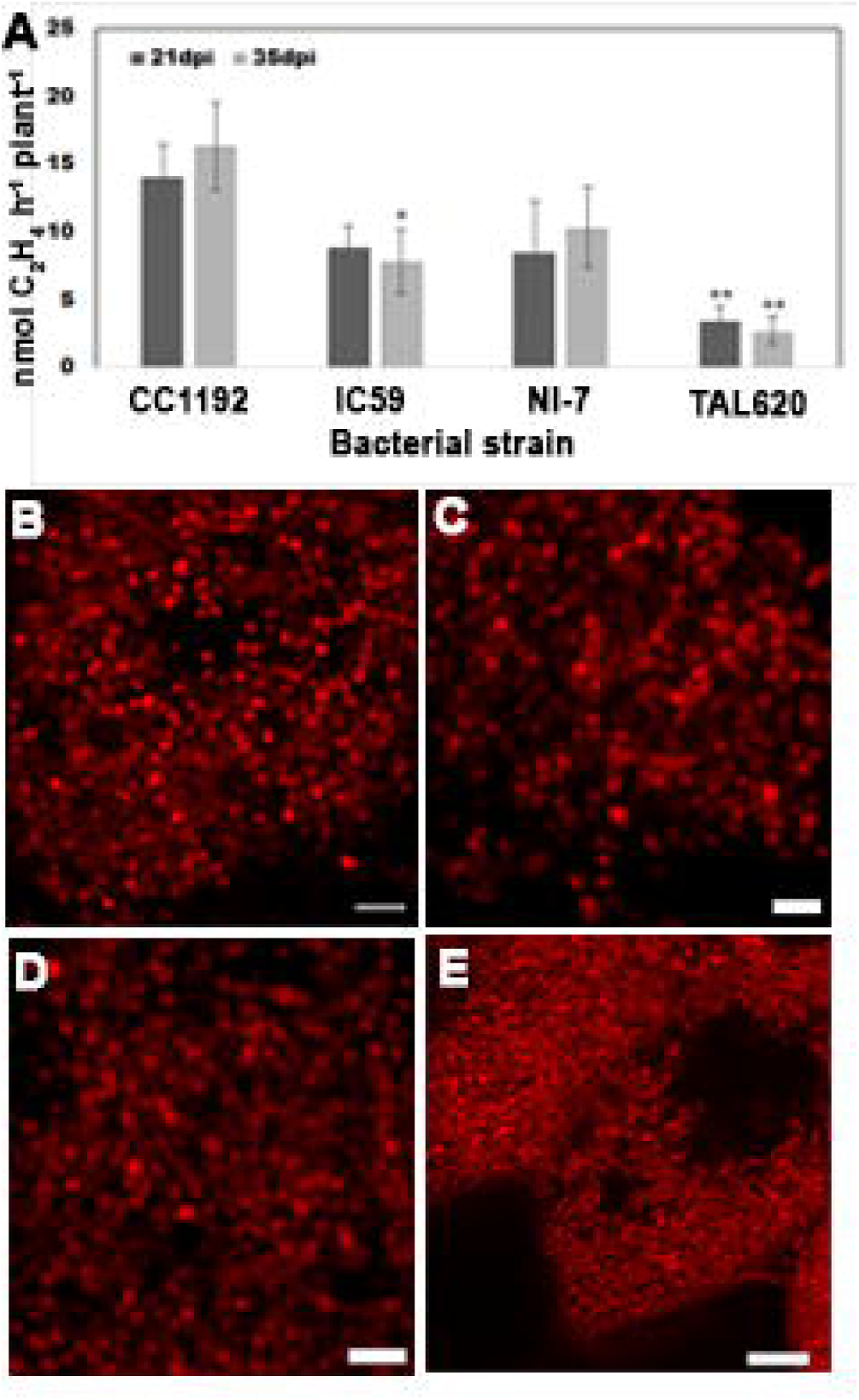
Characterization of different chickpea nodules: **A**, Acetylene reduction activity (ARA) with *Mesorhizobium ciceri CC1192, Mesorhizobium ciceri IC59, Mesorhizobium sp. NI-7* and *Mesorhizobium ciceri TAL620* induced nodules at 21 dpi and 35 dpi. ARA was measured per plant. Mean and SEM of five biological replicates are presented in each case. **B-D**, 21 dpi chickpea nodules were hand sliced and stained with SYTO82 and **E**, SYTO13. **B**, *Mesorhizobium ciceri CC1192* **C**, *Mesorhizobium ciceri IC59* **D**, *Mesorhizobium sp. NI-7*. **E**, *Mesorhizobium ciceri TAL 620* induced nodules in *BDG256* chickpea ecotype. Scale bars in **B-E**, = 5µm. Asterisks indicate significantly less acetylene reduction by *IC59* and *TAL620* induced nodules: *P ≤ 0.05, **P ≤ 0.01

**Figure 4:**
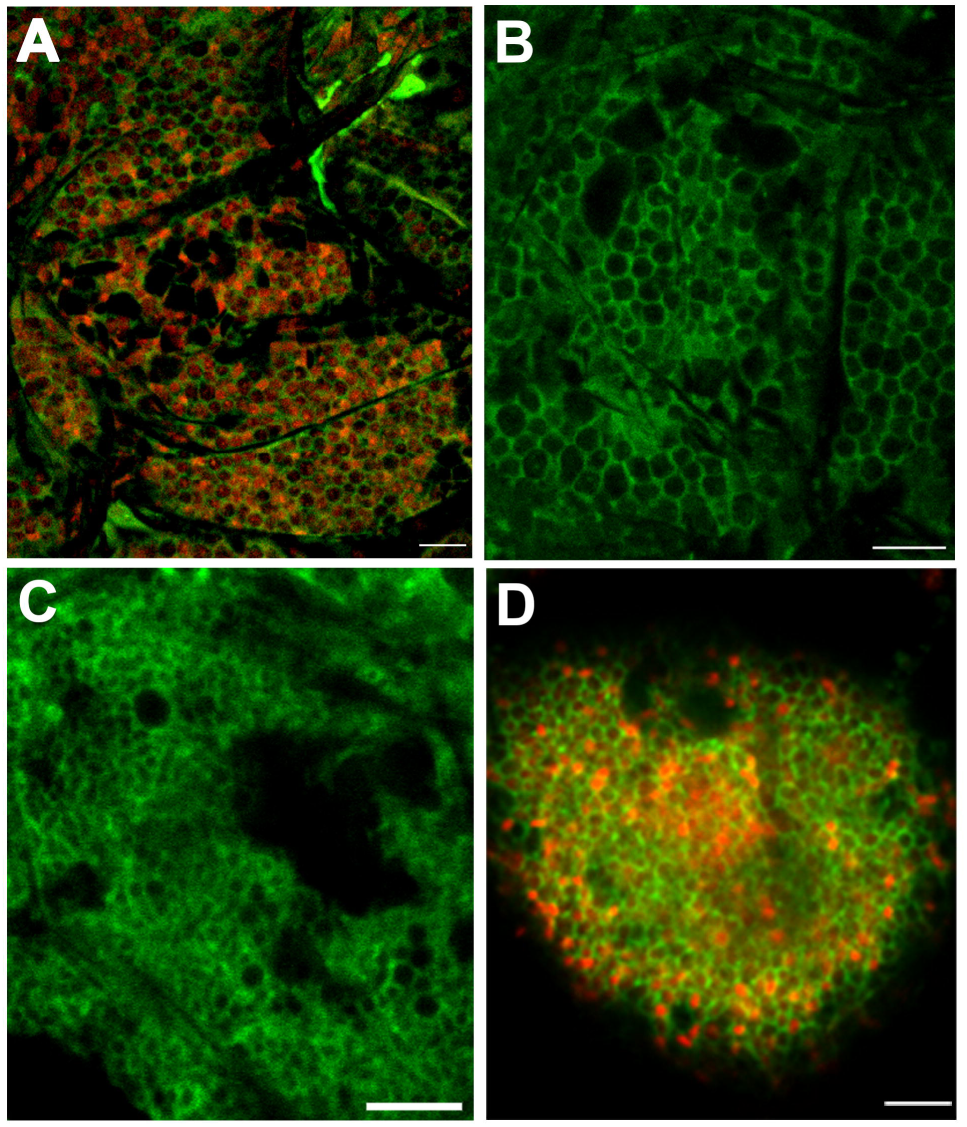
Characterization of the acidity of the peribacteroid space of chickpea nodules: **A-D**, 21 dpi chickpea nodule were hand sliced and stained with DND189 (Acidic compartment, green). **A**, Counterstain with SYTO82 (DNA, a live cell). **D**, Counterstained with PI (DNA of a dead cell). **A-B**, Nodules were induced by *Mesorhizobium ciceri CC1192*. **C-D**, Nodules were induced by *Mesorhizobium ciceri TAL 620*. Scale bars in **A-C**, = 5µm and **D**, = 10 µm.

## Discussion

Unlike model legumes, nodule development studies in chickpea are limited. Nodule development in chickpea and *Medicago* follow a similar course. Hence, the knowledge on *Medicago* nodule development holds true for many aspects of nodule development in chickpea. Nonetheless, differentiation of bacteroids in both the systems show major dissimilarity. Bacteroid differentiation takes place in the invasion zone of both *Medicago* and chickpea nodules. Bacteroids inside *Medicago* nodules endoreduplicate up to 64 fold, whereas a maximum of 4 fold endo-reduplication is observed in chickpea (Montiel et al., 2017). Certain *NCR* genes (*dnf4-NCR211* and *dnf7-NCR169*) are indispensable for nitrogen fixation in *Medicago* (Horvath et al., 2015; Kim et al., 2015). Recently, it has been shown that two *NCR*, nitrogen fixation specificity (*NFS1* and 2) genes also function as negative regulator of symbiont persistence (Wang et al., 2017; Yang et al., 2017). Such a specific role of few NCR peptides amidst overall redundancy among the members of this family suggests complex regulation behind the evolution of terminal differentiation of bacteroids. Also, the presence of a much lower number of *NCR* genes and absence of the symbiotically essential *NCRs* in chickpea genome suggest major differences in the molecular regulation of bacteroid differentiation (Montiel et al., 2017).

The recalcitrance of chickpea towards transformation and it’s low regeneration efficiency is a bottleneck at this moment (Indurker et al., 2007; Varshney et al., 2010); Hairy root transformation is an easy alternative to stable lines, for root biologists. The co-transformed roots can be used for over-expression, knockdown or *CRISPR/Cas9-*mediated deletion of a gene of interest (Limpens et al., 2009; Ron et al., 2014). Till date, the majority of literature on RNS in model legume involves hairy root transformation. The transformation efficiency of model legume ranges around 60-70%. On the other hand, the transformation efficiency of the large-seeded grain legumes soybean, *Vigna,* and pea varies greatly between 20-80% (Sinharoy et al., 2015). According to our protocol, the transformation efficiency of chickpea varies greatly between ecotypes, ranging between 26-53% (Table 1). This is comparable to other large-seeded grain legumes. These transformed roots develop healthy and active nodules (Figure 1) which can fix nitrogen efficiently (data not shown). We have tested our protocol using nine different constructs carrying different transgenes, and all of them showing an expected pattern and subcellular localization (Figure 2 and Supplemental Figure 1). It should be noted that the transformation frequency using *pCMU-ACTLr* (actin cytoskeleton) construct was very low (data not shown).

Our data show that peroxisomes are enriched around the vascular bundle in the nodule cortex (Figure 2). Peroxisomes play a major role in ureide transport inside the determinant nodules of soybean (Hanks et al., 1981). The role of peroxisomes in amide transporting nodules of temperate legumes is not clear. Scavenging of toxic H2O2 around the nodule cortex could be one possibility. Presence of active peroxisomes has been reported also in the meristem and the invasion zones of pea nodules (Borucki, 2007). On contrary, we have noticed a higher number of peroxisomes in the nodule cortex (Figure 2A). Discrimination between active and inactive peroxisomes was beyond the limits of the present study. Our observation demands further studies on the significance of peroxisome accumulation around the nodule cortex of temperate legumes like chickpea. In *Medicago*, the ER apparatus is assembled within the epidermal cells around an area where the future infection thread would penetrate, even before the actual penetrance (Genre et al., 2005). The localization of ER around the infection threads in chickpea nodules indicates similar organization (Figure 2F). ER-medicated trafficking is important during endocytosis of bacteroids. Normally, proteins of exocytosis pathway go from ER to the membrane, through the golgi bodies (Ivanov et al., 2012; Gavrin et al., 2017). Surprisingly, we did not find golgi bodies around the infection threads. The significance of selective concentration of ERs in combination with the absence of golgi bodies around the infection threads is not clear.

The efficiency of nitrogen fixation differs significantly depending on bacterial genotypes. A systematic effort to evaluate the behavior of different *Mesorhizobium* strains during chickpea nodule development has not been given till date. Our study highlights a striking difference of symbiotic performances between different *Mesorhizobium* strains (Figure 3 and Supplemental Figure 2). Presence of strain-specific negatively regulating *NCR* gene in *Medicago* genome (Wang et al., 2017; Yang et al., 2017) opens up the question that is there any *NCR* gene in chickpea genome that causes toxic effect in *TAL620* induces nodule? We detected sporadic spherical-shaped dead bacteroids inside *TAL620* induced nodules (Figure 4D). This, together with the late onset of pink coloration and early senescence of the above-mentioned nodules indicate a delay in bacteriod differentiation and less survival time. 63 *NCRs* express in chickpea *BDG256* nodules induced by *TAL620*, as shown by transcriptomic experiments (Kant et al., 2016; Montiel et al., 2017). This suggests that combinations of at least 63 NCRs are required for the development of this uniquely-shaped symbiosome. Our study highlights the importance of transcriptomic study needs to be performed on chickpea nodules containing normal spherical-shaped symbiosomes. A comparative transcriptomic study between nodules containing spherical and elliptical symbiosomes will highlight the *NCR*s behind the transition between these two unique shapes. It is also very important to screen diverse chickpea ecotypes inoculated by both *TAL620* and *CC1192*/*NI-7* to see whether different hosts can generate different morphotypes of bacteroids.

Overall, our studies with different bacterial strains along with the efficient hairy root transformation protocol will render the community of nodule biologists a new platform on which genetic experiments of chickpea could easily be performed. Moreover, we have presented enough intriguing features of nodule development in chickpea, which merits further attention in future.

## MATERIALS AND METHODS

### Plant material and growth condition

Seeds of chickpea (*Cicer arietinum*) varieties *ICC4958, BDG256, ICC17258, ICC1885, ICC8261* and local varieties (Desi) were surface sterilized by 30% commercial bleach solution (RIN, Hindustan Unilever Limited) containing active sodium hypochlorite 1.2% for 3 minutes. The seeds were thoroughly washed with sterile water. They were placed on plates containing sterile wet filter paper for germination. The plates were kept in 20-22ºC temperature in dark. After 2-3 days, the germinated seedlings were placed in pots with growing substrate (A mixture of 3:1 vermiculite and fire clay balls). The pots were placed in a growth chamber under 200 µE m^−2^ s^−1^ for 16 hours light and 8 hours dark cycle, at 22ºC and 40-60% relative humidity.

### Bacterial material and growth condition

The *Agrobacterium rhizogenes R1000, ARqua1* & *MSU440* strains were grown at 28ºC in solidified Luria Bertani medium with appropriate antibiotics. The *Agrobacterium* strains were transformed with the vectors (*pCMU-ERr, pCMU-APr, pCMU-ACTLr, pCMU-MITr, pCMU-PERr, pCMU-GAr, pCMU-PDESr, pm-ck CFP CD3 1001, pDR5-GFP-NLS* and *pKGW Red root* (Karimi et al., 2002; Nelson et al., 2007; Suzaki et al., 2013; Ivanov and Harrison, 2014) by the freeze-thaw method (Hofgen and Willmitzer, 1988). For hairy root transformation, *Agrobacterium* was grown for 48 hours at 30ºC in LB agar plate with suitable antibiotics to produce a lawn in the plate. *M. ciceri IC59, M. ciceri TAL 620, M. ciceri CC1192* and *Mesorhizobium sp. NI-7* were cultivated at 28ºC in Yeast Mannitol broth.

### Hairy root transformation

To conduct the *in-vivo* method of transformation, seeds were surface sterilized and germinated. The growing tender roots were cut (1-3cm) below epicotyls close to the cotyledonary region, 2-4 days after germination, with a sterile blade. The wounded region of the cotyledon was rubbed in *Agrobacterium* lawn with the appropriate vectors. Alternatively, the germinated seedlings were pierced by a needle containing *Agrobacterium* in the tip. The needle was pricked in epicotyls close to the cotyledonary region of the root. The infected seeds were transferred to a plate containing solidified Fahraeus medium (Boisson-Dernier et al., 2001) without antibiotics. The lower part of the plate was covered with Aluminum foil. The plates were placed longitudinally allowing the root to grow gravitropically, at 22ºC under 16h/8h light and dark cycle. After 7 days hairy root production was observed. When the roots were coming from the injection site, the main roots were cut. Transformed roots were screened under either the Leica stereo fluorescence microscope M205FA equipped with a Leica DFC310FX digital camera (Leica Microsystems) or stereo fluorescence microscope Nikon AZ100 equipped with Nikon digital camera (Nikon Digital Sight DS-Ri1) (Nikon). Non-transformed roots were removed by cutting the roots below the injection region. After 14-15 days the composite plants were transferred to a solidified Fahraeus medium with suitable antibiotics in culture tubes (25×150mm). After 12-15 days the seeds were transferred to growing substrate. Alternatively, the plants with transformed roots were directly transferred in growing substrate without any prior antibiotic selection. The growing substrate was pretreated with full nitrogen B and D medium (Broughton and Dilworth, 1971). The plants were bottom watered four times per week with sterile water. After 10 days in both cases, the plants were treated with full nitrogen B and D medium. After 10-15 days the growing substrate containing plants were washed thoroughly with distilled water and transferred to a new pot for removal of nitrogen. For 2-3 days the plants were maintained in this nitrogen free condition and after that, the plants were inoculated with *Mesorhizobium* for nodulation.

For *ex-vitro* transformation, plants were grown in pots in 22ºC. A slanted cut (5-6cm) was made in the young branches of 25-30 days grown plants. The wounded region was dipped in a bacterial culture plate (Containing 10mM MES pH 5.6 and 20µM acetosyringone) of *Agrobacterium* strains with the appropriate vector. Infected segments were transferred into 4 cm long pots with growing substrate. The pots were placed in a tray containing sufficient full nitrogen B and D medium to overcome water stress. A humid chamber was created around the tray with clear wrap and placed in a growth chamber under 16h/8h light and dark cycle at 22ºC. Plants were regularly checked for maintaining water and humidity. After 15 days roots were seen and after 30 days the growing substrate containing plants were washed thoroughly with distilled water and transferred to a new pot for removal of nitrogen. After maintaining this nitrogen free condition for 2-3 days the plants were inoculated with *Mesorhizobium* for nodulation.

### Nodulation Assay

The growing substrate was vigorously washed for removal of nitrogen prior to nodulation. It was tested for ammoniacal nitrogen and nitrate nitrogen with the soil testing kit (Himedia). The inoculum was prepared from different *Mesorhizobium* culture of OD_600_ (0.7-0.9) in Yeast Mannitol medium. The bacterial culture was harvested, media was discarded and diluted in nitrogen-free B and D media. Each pot was inoculated with 50 ml of diluted 0.03-0.05 OD_600_ bacterial culture in B and D medium.

### Acetylene Reduction Assay

Acetylene reduction assay was performed as described previously (Oke and Long, 1999). Briefly, plants were grown on growing substrate and inoculated separately with the four respective bacteria. At 21 dpi and 35 dpi plants were harvested (5 biological replicates of plants inoculated with each bacterium in each time point). Entire root systems were placed in a test tube and seal with suba-seal septa (Sigma-Aldrich) (1 plant per tube). The tubes were kept in dark at room temperature for 16 hours. Ethylene production was measured by gas chromatography (Shimadzu GC-2010 equipped with HP-PLOT ‘S’ Al_2_O_3_ 50m, 0.53 mm column (Agilent technologies). The mean and standard error of mean (SEM) were calculated for five biological replicates.

### Confocal microscopy

The plants were harvested and the sample was hand sectioned and mounted with water. The staining was done by SYTO13 (DNA, Green) (Invitrogen), SYTO82 (DNA, red) (Invitrogen), Propidium iodide (PI) (DNA of dead cell, red) or DND189 (Acidic compartment, Green) (Invitrogen). The confocal microscopy was performed either in Olympus Model IX81 or in Leica TCS SP5 confocal microscope using an excitation wavelength and emission bandpass of 488 nm and 500-530 nm for SYTO13, 541 nm and 550-580 nm for SYTO82, 448 and 500-530 nm for DND189, 587 nm 600-630 nm for mCherry, 535 nm and 610-640 nm for PI.

### Isolation of *Mesorhizobium* from chickpea nodules

The plants were collected from NIPGR experimental field. Healthy looking pink nodule containing plants were selected. Roots were washed thoroughly to remove soil. About 10 pink nodules were collected by cutting the nodules from the root about 0.5 cm on each side. The nodules were placed in sterile water with 0.12g glass beads and vortexed for 30-60 seconds in repetitive cycles to remove adhering soil particles. It was treated with 30% commercial bleach solution (RIN, Hindustan Unilever Limited) containing active sodium hypochlorite 1.2% for 3 minutes and washed with sterile water for 5-6 times. After surface sterilization, the nodules were rolled over solidified Yeast Mannitol medium and incubated for 2 to 3 days at 28ºC (control). Immediately, the same nodules were squashed in sterile water. It was serially diluted with sterile water and plated in solidified Yeast Mannitol medium. If the control plate does not shows growth, then only the corresponding experimental plates were processed for further characterization. The experimental plates were placed in 28ºC for 5 to 6 days. The growth was checked regularly. The bacterial colony in the experimental plates were subculture to purify the bacterial culture and further used for the nodule developmental experiment.

### Cloning of 16S rDNA

*NI-*7 *16S rDNA* was amplified by forward 5′ TAACACATGCAAGTCGAACG 3′ and reverse 5′ ACGGGCGGTGTGTAC 3′ primers using PrimeSTAR Max DNA polymerase (Takara, Clontec). It was then cloned in pJET 1.2/blunt vector (Clone JET PCR cloning kit, Thermo Fisher). Sanger sequencing was done with sequencing primers provided by Clone JET PCR cloning kit (forward 5′ CGACTCACTATAGGGAGAGCGGC 3′ and reverse 5′ AAGAACATCGATTTTCCATGGCAG 3′). The sequence was assembled and processed with Geneious v9.1.8. The *16S rDNA* sequence of the *NI-7* strain has been submitted to NCBI (MH517450)

**Supplemental Figure 1:** Localization of different subcellular protein markers in chickpea hairy roots: **A**, mCherry-HDEL localization to the ER. **B**, COX4-mCherry localization to mitochondria **C**, LifeAct-mCherry localization to actin microfilaments, **D**, mCherry-SKL localization to peroxisome **E**, BCPsp-mCherry localization to apoplastic space **F**, PIP2A-CFP localization to the plasma membrane in the chickpea epidermal cells. **G**, PDLP1-mCherry localization to plasmodesmata in the chickpea cortical cells **H**, MAN49-mCherry localization to golgi apparatus (cis-golgi) in the chickpea epidermal cells **I**, Localization of GFP-NLS to the nucleus of chickpea root primordia driven by the *DR5* promoter. Nucleus marked by N and vascular bundle marked by V. Scale bars in **B-D**, **G**, = 10µm, **A**, **E-F**, **H**, = 5µm and **I**, = 50µm.

**Supplemental Figure 2:** Nodule developmental time course of chickpea nodules: **A-D**, 6 dpi nodules. **E-H**, 10 dpi nodules **I-L**, 15 dpi nodules **M-P**, 21 dpi nodules **Q-T**, 28 dpi nodules **U-X**, 35 dpi nodules **A**, **E**, **I**, **M**, **Q**, **U**, *Mesorhizobium ciceri CC1192* induced nodules **B**, **F**, **J**, **N**, **R**, **V**, *Mesorhizobium ciceri IC59* induced nodules **C**, **G**, **K**, **O**, **S**, **W**, *Mesorhizobium* sp. *NI-7* induced nodules **D**, **H**, **L**, **P**, **T**, **X**, *Mesorhizobium ciceri TAL 620* induced nodules. Scale bars **A-X**, = 1mm.

## Acknowledgement

We thank R. Varshney and S. Gopalakrishnan, ICRISAT, India; S. Bhatia, NIPGR-New Delhi, J. Terpolilli, Murdoch University, Australia for providing *M. ciceri IC59, M. ciceri TAL 620*, and *M. ciceri CC1192* respectively; H.D. Upadhyaya and D. Sastry, ICRISAT, India for providing chickpea seeds (*ICC4958, ICC17258, ICC1882, ICC8261*); S. Bhatia, NIPGR for providing *BDG256*. D. J. Chattopadhyay, Amity University; M. DasGupta of Department of Biochemistry, University of Calcutta, and A. Seal, Department of Biotechnology, University of Calcutta, for their enormous support and allowing us to use their facility. A. Seal for providing *pm-ck CFP CD3 1001* construct, Maria J. Harrison, Boyce Thompson Institute for Plant Research Ithaca, USA for providing pAtUb driven subcellular marker constructs and Suzaki Takuya, University of Tsukuba, Japan for providing *DR5-GFP-NLS* construct. DBT-CU-IPLS and NIPGR for their confocal facilities; CIF-NIPGR; NIPGR-DELCON for their support. T Khanna, Jamia Millia Islamia, New Delhi for technical support. This work was supported by core research grant from National Institute of Plant Genome Research and Ramalingwaswami Re-entry grant, DBT (BT/RLF/Re-entry/41/2013).

